# Emotional memories are enhanced when reactivated in slow-wave sleep but impaired in REM

**DOI:** 10.1101/2023.03.01.530661

**Authors:** Cagri Yuksel, Dan Denis, James Coleman, Boyu Ren, Angela Oh, Roy Cox, Alexandra Morgan, Erina Sato, Robert Stickgold

## Abstract

Sleep supports memory consolidation. However, it is not completely clear how different sleep stages contribute to this process. While rapid eye movement sleep (REM) has traditionally been implicated in the processing of emotionally charged material, recent studies indicate a role for slow wave sleep (SWS) in strengthening emotional memories. Here, to directly examine which sleep stage is primarily involved in emotional memory consolidation, we used targeted memory reactivation (TMR) in REM and SWS during a daytime nap. Contrary to our hypothesis, reactivation of emotional stimuli during REM led to impaired memory. Consistent with this, REM% was correlated with worse recall in the group that took a nap without TMR. Meanwhile, cueing benefit in SWS was strongly correlated with the product of times spent in REM and SWS (SWS-REM product), and reactivation significantly enhanced memory in those with high SWS-REM product. Surprisingly, SWS-REM product was associated with better memory for reactivated items and poorer memory for non-reactivated items, suggesting that sleep both preserved and eliminated emotional memories, depending on whether they were reactivated. Notably, the emotional valence of cued items modulated both sleep spindles and delta/theta power. Finally, we found that emotional memories benefited from TMR more than did neutral ones. Our results suggest that emotional memories decay during REM, unless they are reactivated during prior SWS. Furthermore, we show that active forgetting complements memory consolidation, and both take place across SWS and REM. In addition, our findings expand upon recent evidence indicating a link between sleep spindles and emotional processing.

## 1. INTRODUCTION

Sleep supports memory consolidation, which allows the stabilization and integration of recently formed labile memories (Stickgold, 2005). However, it is unclear how different sleep stages, with their distinct physiology, contribute to this process. For declarative memories, the predominant model, the active system consolidation theory, posits that memory consolidation is achieved by reactivation of recently encoded, hippocampus-dependent memories during slow-wave sleep (SWS) when the coordinated activity of hippocampal ripples, thalamocortical spindles and cortical slow oscillations distribute these memories to long-term stores in neocortical networks (Klinzing et al., 2019). While this framework is constructed around non-REM sleep (NREM) physiology, and SWS in particular, there is strong evidence that rapid eye movement sleep (REM) is also involved in the consolidation of declarative memories (Boyce et al., 2017). Instead of a NREM-REM dichotomy, alternative but compatible accounts, such as the sequential hypothesis (Giuditta, 2014) and others (Singh et al., 2022), attribute the memory benefit of sleep to successive NREM-REM episodes.

Processing of emotional material during sleep, including emotional memory consolidation, has been traditionally attributed to REM (Davidson et al., 2021). Some studies supported this view, and showed higher retention of emotional content following REM-rich late night sleep but not after early sleep (Groch et al., 2013) or in SWS deprived participants but not in those deprived of REM (Wiesner et al., 2015). In addition, REM measures were correlated with increased retention of emotional memories (Nishida et al., 2009). Based on these findings, it was hypothesized that REM provides a unique milieu which facilitates the consolidation of memory of emotional experiences (Goldstein and Walker, 2014). However, this hypothesis is challenged by an accumulating number of studies which found no association of REM measures with emotional memory performance (Baran et al., 2012; Kaestner et al., 2013; Cairney et al., 2014a; Ackermann et al., 2015; Alger et al., 2018; Sopp et al., 2018; see Davidson et al., 2021 for a review). More importantly, increasing evidence suggests that NREM is involved in emotional memory consolidation. Two studies found NREM to be sufficient for emotional memory consolidation, regardless of the presence of REM (Morgenthaler et al., 2014; Cellini et al., 2016). Furthermore, several studies showed correlations of emotional memory benefit with time spent in NREM (Wagner et al., 2007; Cairney et al., 2015; Payne et al., 2015; Alger et al., 2018) and spectral power in the delta band (Payne et al., 2015). Beyond correlations, pharmacologically increasing slow wave activity (Benedict et al., 2009) and sleep spindle density (Kaestner et al., 2013) enhances memory for emotional items.

Another line of direct evidence for the role of NREM in emotional memory consolidation comes from targeted memory reactivation (TMR) studies. TMR relies on replaying cues that are associated with recently encoded memories and has been shown to enhance declarative memory retention (Hu et al., 2020). In one TMR study, emotional and neutral memories were cued during SWS (Cairney et al., 2014b). While there were no memory benefits, SWS duration and number of spindles were associated with shorter reaction times for emotional items. In another study, which cued emotional and neutral items in NREM or REM, memory for emotional items was improved after cueing in NREM but not REM (Lehmann et al., 2016). Furthermore, successful cueing was associated with increases in spindle and theta power. However, another study found no effect of TMR in NREM on emotional memory (Ashton et al., 2018).

In this study, we used TMR to examine which sleep stage is primarily involved in emotional memory consolidation. Consistent with the prevailing view in the literature on the role of REM in emotional memory consolidation, we hypothesized that replay of cues for emotional items during REM would lead to enhanced retention compared to replay in SWS. We also explored whether emotional memories are retained better than neutral memories. We hypothesized that replay of cues for emotional items would lead to greater memory improvements than the cueing of neutral items.

## 2. METHODS

### 2.A Participants

The study sample included healthy young participants recruited from colleges in the greater Boston area. A total of 185 participants consented to the experiment, of which 123 provided usable datasets for analysis (age: 21.6±2.7; 63.9% female; see below for details). Participants reported no abnormal sleep patterns, history of psychiatric or neurological disorders, or current medication use. They were instructed to keep a regular sleep schedule for the 3 nights preceding the experiment and were asked to refrain from recreational drugs and alcohol for 48 hours, and caffeine in the morning, before their visits. All participants provided written consent approved by the Institutional Review Board of Beth Israel Deaconess Medical Center.

Participants were randomly assigned to one of 5 groups. Emotional SWS (E-SWS) and emotional REM (E-REM) groups learned emotional items and were then exposed to reactivations during SWS or REM, respectively. Similarly, a neutral SWS (N-SWS) group learned neutral items with reactivations in SWS. We also included emotional nap (E-Nap) and emotional wake (E-Wake) groups, which learned emotional items and took a nap without any reactivations or did not take a nap but rested quietly for a similar period before testing. Reactivation groups were matched in age (F=.54, p=.71) and sex (*X^2^*=2.4, p=.66).

### 2.B Design and Procedures

The protocol included 2 visits (**Figure 1.A**). On the first visit, participants arrived at the laboratory at around 11:00 AM. After consenting, they filled out the Epworth Sleepiness Scale (Johns, 1991) and questionnaires about their sleep patterns and quality in the preceding 3 days. They were then wired for EEG (detailed below) and a brief (7 minutes) resting state EEG was recorded with eyes closed. We do not report EEG data from the rest periods in this paper.

**Figure 1.**
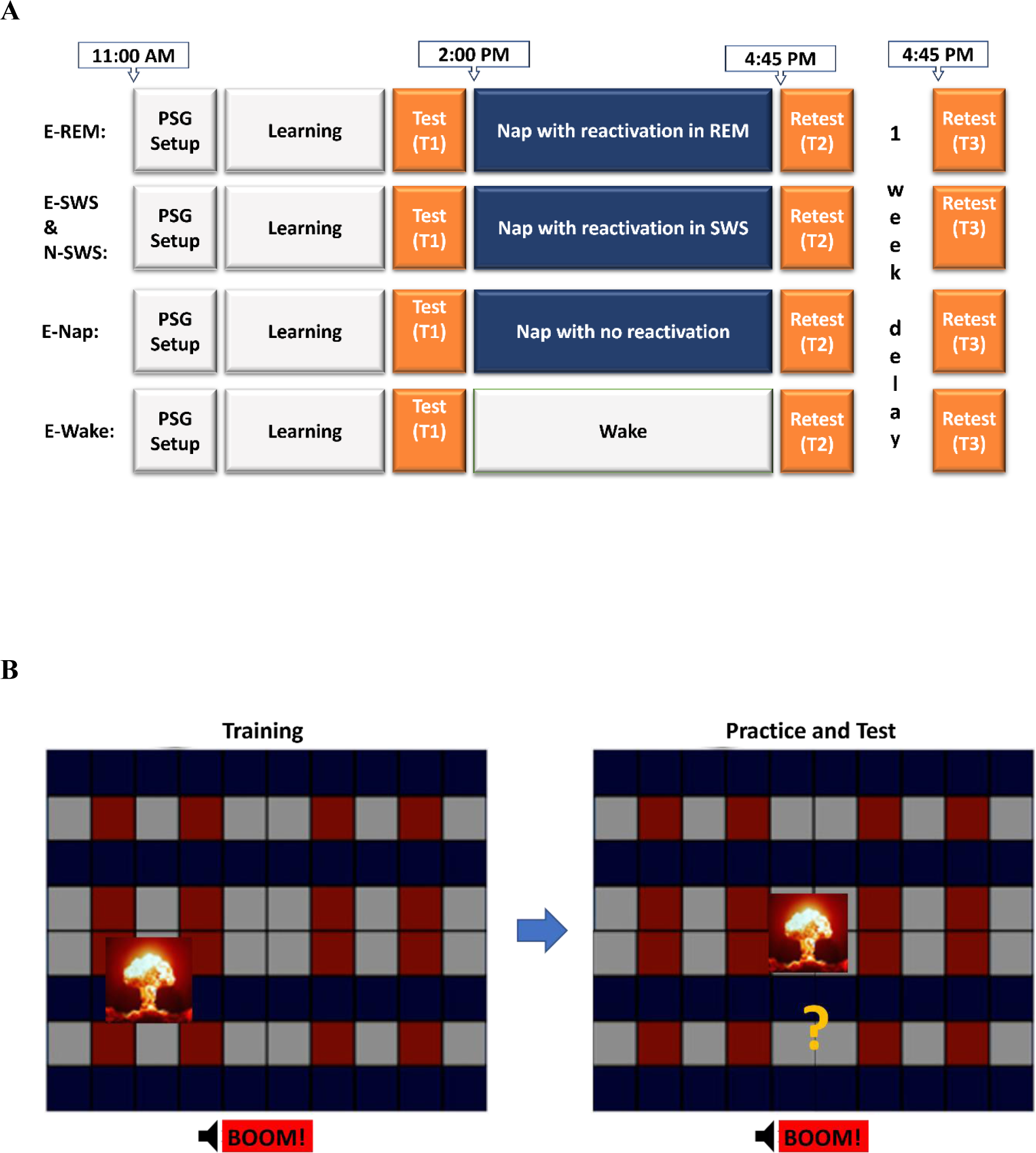
**A** - Timeline of the study procedures for different groups are depicted. In all groups, the first visit started at 11:00 AM. After training and practice, a baseline test (T1) was carried out immediately before the nap. The first retest (T2) was approximately 45 minutes after waking up. The second retest (T3) was approximately 1 week after the first visit. **B –** During training, participants passively viewed emotional or neutral pictures appearing at different locations on a grid, while a paired sound was played. During practice and test, pictures appeared in the center of the screen, and participants were asked to move them to their correct location.

We used a modified version of the TMR task described in previous publications (Rudoy et al., 2009; Creery et al., 2015; **Figure 1.B**). Learning included two successive phases, training and practice. During training, participants viewed 50 neutral or negative pictures appearing in different locations on a grid, in random order, while simultaneously hearing a 1-second sound that was naturally linked with that object. During practice, pictures appeared in the center of the screen while their corresponding sound played. Participants were instructed to move the objects to their original location and press the mouse button. After the mouse was clicked, the picture was moved to its correct location, which provided feedback. In the first two runs of the practice phase, all 50 objects were tested. In subsequent rounds, items that had been placed within 150 pixels of their correct locations in two successive rounds no longer appeared. The practice phase continued until all objects had been removed from the testing pool. The baseline test (T1) followed immediately after learning. This phase was similar to practice, except participants placed each picture only once and no feedback was provided. Following test 1, another 7-minutes resting state EEG was collected.

For participants in the sleep groups, lights were turned off shortly after the baseline test, usually around 2 PM. Sleep was allowed up to 2 hours. For all sleep participants, white noise was played through bedside speakers (∼39 dB on the pillow) starting at lights off and continuing until lights on. In the reactivation groups, half of the sounds (n=25) were presented in a random order, with 5-second interstimulus intervals. Sound presentations began approximately 1.5 minutes after SWS or REM onset. Replayed sounds were selected by a computer algorithm so that memory accuracy at T1 for the replayed and non-replayed sounds were similar. Sounds were presented until the specific sleep stage (SWS or REM) ended. If the participant entered the same sleep stage again, reactivation was resumed. Data from a participant were included in analyses if all 25 sounds were played at least once. After lights on, a final resting state EEG data was collected. Wake participants were also wired for EEG and spent an equal amount of time in the bedroom, doing relaxing activities such as reading, while they were observed to ensure wakefulness.

Retests were carried out the same way as Test 1. The first retest (T2) took place approximately 45 minutes after lights on, or at the end of the rest period for the wake group. Before they left the laboratory, participants were instructed that their memory would be tested, in the same way, at their second visit. The second visit was approximately one week later and included only the delayed retest (T3), which took place at 4:45 PM, to match the timing of Test 2. EEG was not monitored at this visit.

Before training and each retest, participants filled out the Stanford Sleepiness Scale (Hoddes et al., 1972) and a two-item questionnaire about their ability to concentrate and their level of “feeling refreshed”. After the tests, they were asked to report how difficult or easy and how boring or interesting the tests were, and the strongest emotion they felt.

### 2.C Stimuli

50 emotional and 50 neutral images from online picture databases, the International Affective Pictures System (Lang et al., 2008) and the Geneva Affective Picture Database (Dan-Glauser and Scherer, 2011), and Google image search, were used. Images were 150 x 150 pixels and were displayed on a 67.5 cm x 57.25 cm monitor with a viewing resolution at 1440 x 900 pixels. Emotional images were only negative. 1-second sounds that were naturally linked to each of these images were then taken from the database Pond5 (www.pond5.com) and were paired with each image.

Prior to the main study, a pilot study was carried out to confirm that the emotional and neutral stimulus sets were significantly different in emotion ratings. This study confirmed that both emotional sounds and emotional sound-picture pairs were significantly more negative than their neutral counterparts (See Supplementary material).

### 2.D EEG Acquisition and preprocessing

EEG data was acquired from 57 channels (positioned according to the 10-20 system). Additional electrodes were placed on the left and right mastoids, above the right eye and below the left eye (for EOG), two placed on the chin (for EMG), one on the forehead (recording reference), and one on the collarbone (ground). Data were collected with an Aura-LTM64 amplifier and TWin software (Grass Technologies). All impedances were kept to <25 kOhm. The sampling rate was 400 Hz.

Sleep scoring was performed using TWin software and MATLAB (The MathWorks, Natick, MA)) according to AASM criteria (Berry RB, 2020). Subsequent EEG analyses were performed in MATLAB using custom scripts. First, all EEG channels were re-referenced to the average of the two mastoids, high-pass filtered at 0.3Hz, and notch filtered at 60Hz. Data were then artifact rejected based on visual inspection, with bad segments of data being marked and removed from subsequent analyses. Bad channels were identified by visual inspection and interpolated using a spherical spline algorithm. All artifact-free data were then subjected to further analysis.

### 2.E EEG data analysis

Clean, artifact-free data were segmented into epochs that extended from 1 second before stimulus-onset to 3 seconds after and baseline adjusted to the mean voltage during the 1-second before cue onset. Complex Morlet wavelets were used to decompose the epoched and baseline adjusted time series data into time-frequency representations (Cohen, 2014), with spectral power being extracted at 30 logarithmically spaced frequencies from 2-40Hz and the number of wavelet cycles increasing from 3 to 10 in 30 logarithmically spaced steps to match the number of frequency bins. For analysis, power was decibel-normalized within-subject (10 x log10(power/baseline), where the baseline was mean power in the 200-500 milliseconds prior to cue-onset. This baseline period was chosen to mitigate contamination of the baseline period by post-stimulus activity. As such, positive values reflect relative increases in power following sound cues compared to the pre-cue baseline, whereas negative values reflect decreases. Time-frequency analyses were conducted at electrode site Cz only.

### 2.F Data reduction

Some participants’ data were excluded, because of achieving less than one full round of reactivation (n=32), having a full round of reactivation carried out in the wrong sleep stage (n=4), having too short or fragmented sleep (sleep <45 minutes or WASO>30 minutes; n=18), due to equipment failure (n=5), failing to complete the experiment (n=2), or not meeting healthy control criteria (n=1). In addition, 1 participant from the E-REM group was removed because their error at T1 (also at T2) was more than 3 standard deviations from the mean. The final sample included 24 participants in E-SWS, 25 in E-REM, 31 in N-SWS, 22 in E-Nap, and 20 in E-Wake. Finally, EEG of 1 participant in N-SWS was missing from the spectral analyses.

### 2.G Statistical Analyses

Recall accuracy was measured as the distance between where participants placed the pictures and their correct locations, measured in pixels, with smaller errors in placement indicating better recall accuracy. Because error was not normally distributed within subjects, we used medians in all analyses. The effect of reactivation on recall accuracy within groups was examined using linear mixed-effects models (LMM) with % change in error from T1 to T2 or to T3 (%T1-T2 or %T1-T3, respectively; larger values indicated more forgetting) included as the dependent variable, and reactivation included as a fixed effect. To compare the effect of reactivation between groups, group and (group × reactivation) interaction were included as additional fixed effects. In all models, compound symmetry covariance structure was used for the repeated measures and subject was included as a random effect. A one-way MANOVA was used to compare the sleep stage compositions between groups, with Tukey’s test for post-hoc analyses. Pearson’s correlation was used for correlations between normally distributed variables. Spearman’s correlation was used if variables were not normally distributed.

To analyze cue-evoked EEG data, we first sought to identify clusters of time-frequency points for which post-cue activity was significantly different from zero. For each group, we conducted one-sample t-tests across participants to detect points in the time-frequency space at which spectral power was systematically different from zero (*p* < .05, false discovery rate (FDR) adjusted) (see Schechtman et al., 2021 for a similar approach). Clusters of significant activity were identified using the *bwlabeln* function in MATLAB (Cohen 2014). Within each cluster, spectral power was averaged across all time-frequency points in that cluster, producing for each participant a single spectral power value for each cluster. These values were then used in analyses correlating cue-evoked spectral power with cueing benefit.

## 3. RESULTS

### 3.A Memory Performance

#### Correlation with Sleep Measures

We first examined the correlation of the duration of sleep stages, with cueing benefit (CB; % change in error for the non-reactivated items - % change in error for the reactivated items; (Creery et al., 2015)) and in the case of significant associations, explored correlations with % change in error for reactivated and non-reactivated items separately, to gain further insight into the underlying mechanism for CB.

In the E-SWS group, CB at T2 and T3 (CB-T2 and CB-T3) were positively correlated with %REM (r=.43, p=.03 and r=.44, p=.04, respectively), indicating comparatively less forgetting for reactivated items with increasing proportions of REM. CB-T2 also showed a strong trend towards a significant positive correlation with %SWS (r=.40, p=.051). Because there is evidence suggesting that NREM and REM may have complementary roles in memory consolidation, we also examined the correlation of the product of %SWS and %REM (%SWS × %REM; SWS-REM product) with CB (Stickgold, 2000; Mednick, 2003; Hu, 2015). This revealed a stronger correlation with CB-T2 (spearman’s r_s_=.66, p=4×10^-4^; **Figure 2.A, left**). Considering reactivated and non-reactivated categories separately, to our surprise, higher SWS-REM product was associated with smaller %T1-T2 (less forgetting) for reactivated items (spearman’s r_s_=-.44, p=.03) and with larger %T1-T2 (more forgetting) for non-reactivated items (spearman’s r_s_=.65, p=.001) (**Figure 2.A, right**), suggesting that the combination of SWS and REM facilitated both the strengthening and decay of emotional memories, depending on whether they were reactivated during SWS. Additionally, duration of stage 2 sleep (%N2) was inversely correlated with CB-T2 (r=-.47, p=.02) and positively correlated with %T1-T2 for reactivated items (r=-.47, p=.02), likely reflecting that more %N2 was associated with less SWS-REM product (r=-.49, p=.01).

**Figure 2.**
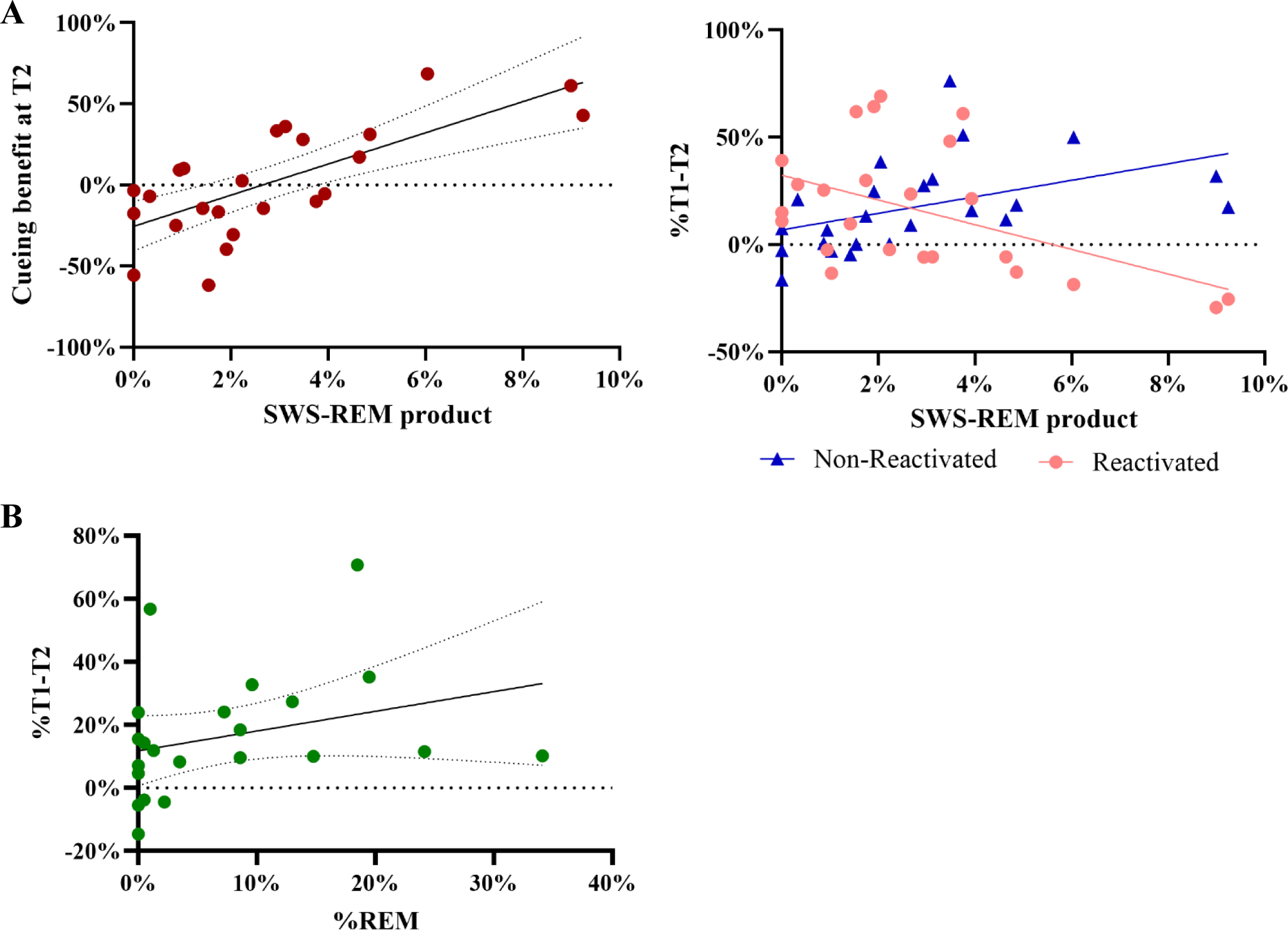
**A** - Correlation of SWS-REM product with CB-T2 (left) and %T1-T2 (right), in E-SWS. **B-** Correlation of %T1-T2 with %REM in E-Nap. Note that larger %T1-T2 indicates more forgetting. Dotted lines indicate 95% confidence interval.

In the E-Nap group, %T1-T2 was positively correlated with %REM (spearman’s r_s_=.46, p=.04; **Figure 2.B**), indicating that REM was associated with memory impairment for emotional items.

#### Effect of TMR on Memory

Next, we examined the effect of reactivation on change in error (%T1-T2 and %T1-T3), in separate groups, using LMMs. For %T1-T2, E-REM showed a significant main effect for reactivation (β[SE] = -.11 [.05], t(24)=2.13, p=.04), and contrary to our expectations, displayed a larger increase in error for the reactivated items (22% for reactivated items vs. 11% for non-reactivated items), indicating that REM cueing was associated with memory impairment, in line with the finding in E-Nap. In E-SWS, for %T1-T2, (reactivation × SWS-REM product) interaction was significant (F = 23.43, p = 8×10^-5^), which aligned with the previously noted correlations in opposite directions for reactivated and non-reactivated items (**Figure 2.A, right**), and indicated that the effect of reactivation was dependent on SWS-REM product. This dependency was reflected in the fact that while there was no significant reactivation effect in the whole group (β[SE] =.017[.07], t(23)=.24, p=.81), those above the median for SWS-REM product (high SWS-REM subgroup) showed a significant benefit for reactivation (β[SE] =.24 [.08], t(11)=3.11, p=.01). In N-SWS, there was no significant main effect of reactivation on %T1-T2 (β[SE]=.03[.05], t(30)=.57, p=.57). For %T1-T3, there was no significant reactivation effect in any of the groups. %T1-T2 and %T1-T3 for all groups and the results of LMMs can be found in supplementary figure S1, and supplementary tables S1 and S2, respectively.

In the LMMs that compared the effect of reactivation on %T1-T2 between groups (E-REM vs. E-SWS, and E-SWS vs. N-SWS), SWS-REM product and its interactions with reactivation and group were included as additional fixed effects, because SWS-REM product was correlated with CB-T2 and the (SWS-REM product × reactivation) was significant in E-SWS (see above). In the analysis that included E-SWS vs. E-REM, there was no (group × reactivation) interaction (β [SE] = .16 [.14], t(45)=1.12, p=.27) **(Figure 3.A; supplementary Table S3)**. On the other hand, in the analysis that included E-SWS with N-SWS, this interaction was significant, indicating that reactivation of emotional items showed a larger memory benefit compared to neutral items (β [SE] = .39[.12], t(49)=3.36, p=.002) **(Figure 3.B; supplementary Table S3)**. In the analyses that compared the effect of reactivation on %T1-T3, %REM and its interactions with reactivation and group were included as additional fixed effects, because it was correlated with CB-T3 (see above). There was no significant (group × reactivation) interaction in either analysis (**Supplementary Table S4)**.

**Figure 3.**
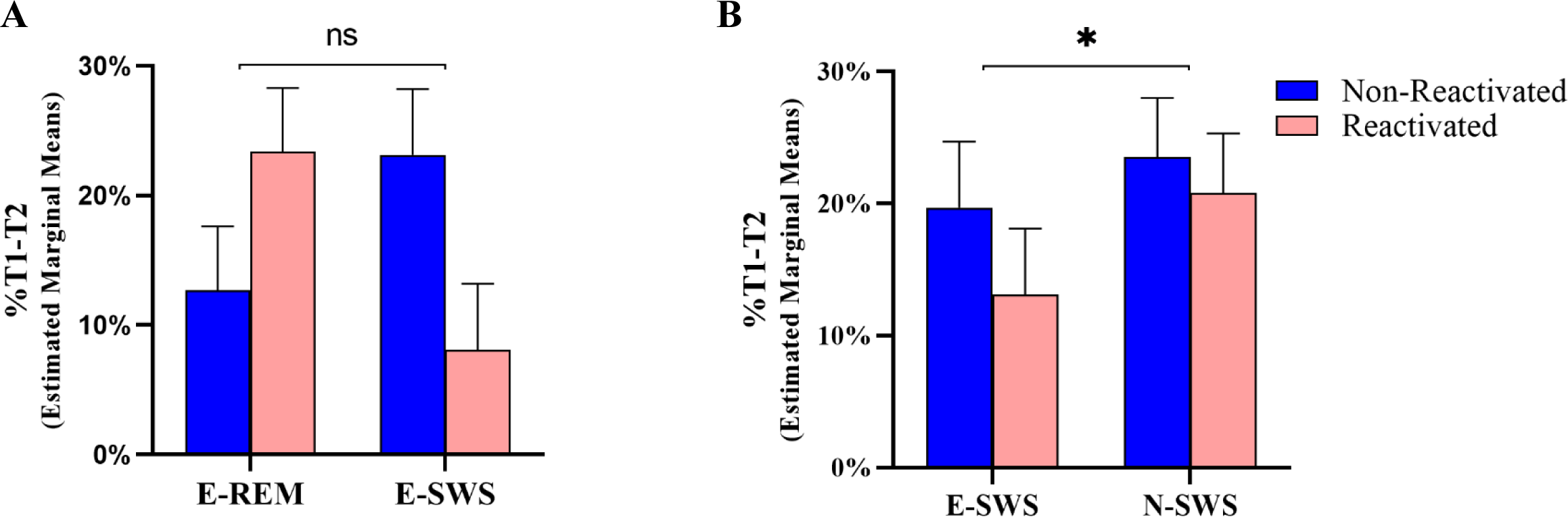
Estimated marginal means for %T1-T2 in linear mixed-effects models that tested the reactivation × group interactions in E-REM and E-SWS (A) and in E-REM and N-SWS (B). Error bars indicate the standard error. *: p<0.05

In summary, emotional memories were impaired when reactivated in REM, and improved when reactivated in SWS, if high SWS-REM product was attained. In addition, reactivating emotional memories in SWS provided a larger memory benefit than reactivating neutral memories.

### 3.B Baseline Memory, Sleep Architecture and Recall in Nap and Wake Groups

As expected, recall immediately after training, at T1, for items that would subsequently be reactivated did not differ significantly from recall of those that would not be reactivated in any of the groups (all p values >.25) **(Supplementary Table S1)**. There was also no difference in recall at T1 between groups (F=1.91, p=.15). Finally, we compared the sleep architecture between groups included in the regression models (E-SWS vs. E-REM; E-SWS vs. N-SWS) and found a higher %REM in E-REM than in E-SWS (p<.001) and N-SWS (p=.01) **(Supplementary Table S2)**. There was no difference in %T1-T2 or %T1-T3, between E-Nap and E-Wake (t(40)=1.44, p=.16 and t(35)=.80, p=.43).

### 3.C Cue-evoked activity

To investigate cue-related modulation of EEG activity during sleep, we first identified clusters of cue-induced activity that were different significantly from zero (see Methods). In the E-SWS group, we identified two time-frequency clusters in which the cues induced increased activity (*p* < .05, FDR adjusted; **Figure 4A)**. The first cluster (2.0 - 8.5Hz), extending from 235-975 ms after cue onset, comprises activity in the canonical delta and theta bands. The second cluster (11.6 - 19.1Hz), occurring 638-1,475 ms following cue onset, corresponds broadly with the sigma (spindle) band. This pattern of increased delta/theta and spindle band activity following the replay of emotional sounds during SWS replicates findings in other studies of cueing during SWS (Lehmann et al., 2016; Forcato et al., 2020; Schechtman et al., 2021).

**Figure 4.**
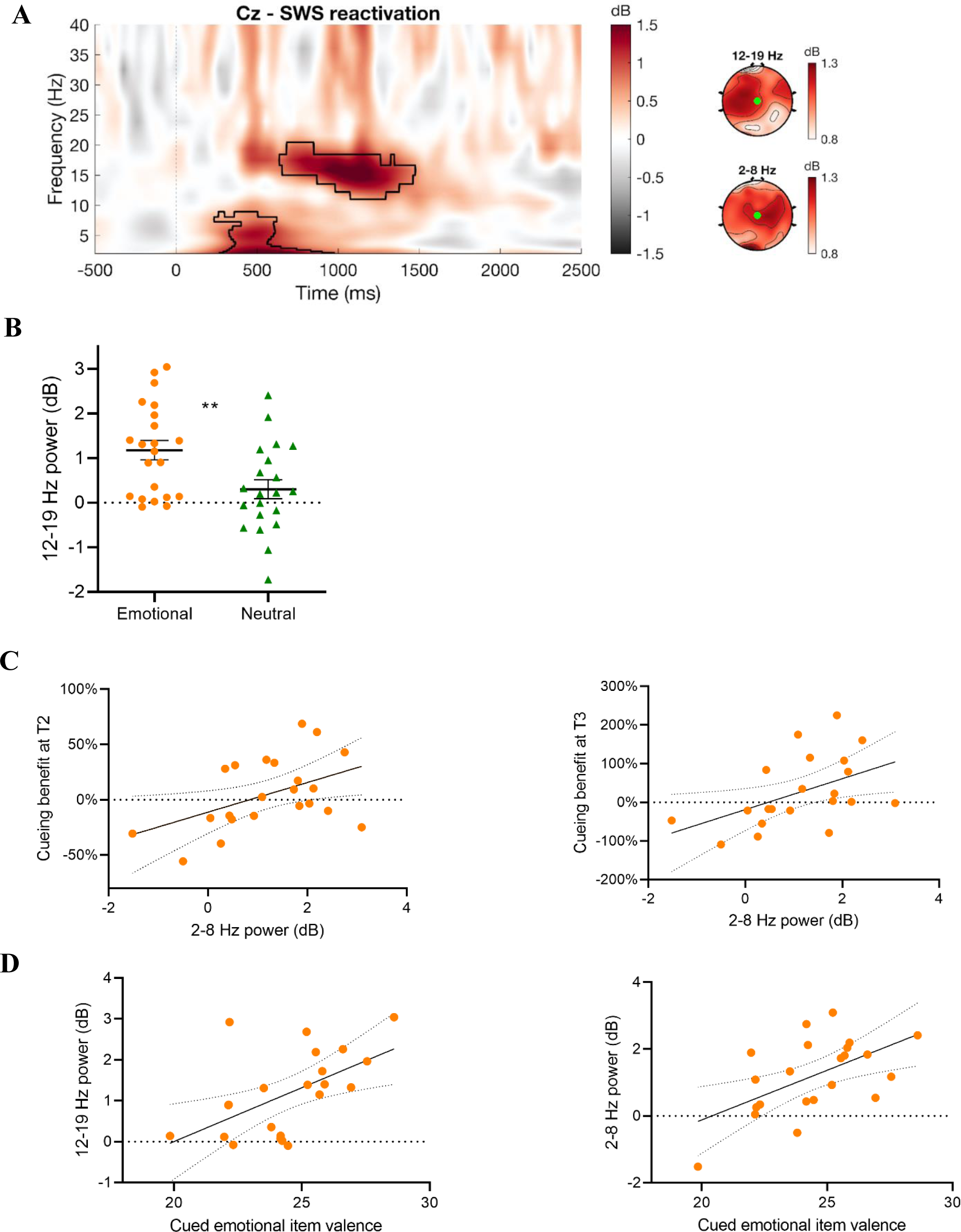
Effect of cueing during slow wave sleep. **A -** Left: Time frequency response to emotional cue presentation at electrode Cz. Time zero represents the initiation of sound presentation during sleep. Black contour lines highlight significant clusters of activation (*p* < .05, FDR adjusted). Right: Topographical visualizations of cluster-averaged activity. **B** - Cluster-averaged time frequency activity in the 12-19 Hz response in response to either emotional or neutral sound cues. Error bars indicate the standard error. ** = *p* < .01. **C -** In E-SWS, 2-8 Hz response is associated with cueing benefit at both T2 (left) and T3 (right). **D -** In E-SWS, valence of the sound cues correlated with both 12-19 Hz and 2-8 Hz responses.

Interestingly, power in the spindle cluster was significantly higher in the E-SWS group compared to the N-SWS group (t(41.9) = 2.99, p = .005; **Figure 4B**), suggesting a difference between emotional and neutral sounds in terms of their modulation of spindle activity. This was further suggested by a significant correlation between the magnitude of the spindle band response and the subjectively rated valence (derived from our pilot study; see supplementary methods) of the emotional sounds cued during sleep (r = .55, p = .008; **Figure 4D, left)**. In E-SWS, power in the delta-theta cluster was positively correlated with the cueing benefit at both T2 (r = .47, p = .03; **Figure 4C, left**) and T3 (r = .48, p = .03; **Figure 4C, right).** The valence of the cued sounds was also correlated with the delta-theta power (r = .56, p = .006; **Figure 4D, right)**.

With regards to cueing during REM sleep, we observed an increase in power across a broad frequency range throughout the entire post-cue period, but the increase was only significant in alpha-beta frequencies (9.4 - 26.5Hz) from 1,375 to 1,725 ms following cue onset (**Figure 5**). Power in the identified cluster did not correlate with the cueing benefit at either T2 or T3 (all p > .43). Similarly, post-cue EEG activity following REM cueing was not related to stimulus valence (r = .12, p = .61).

**Figure 5.**
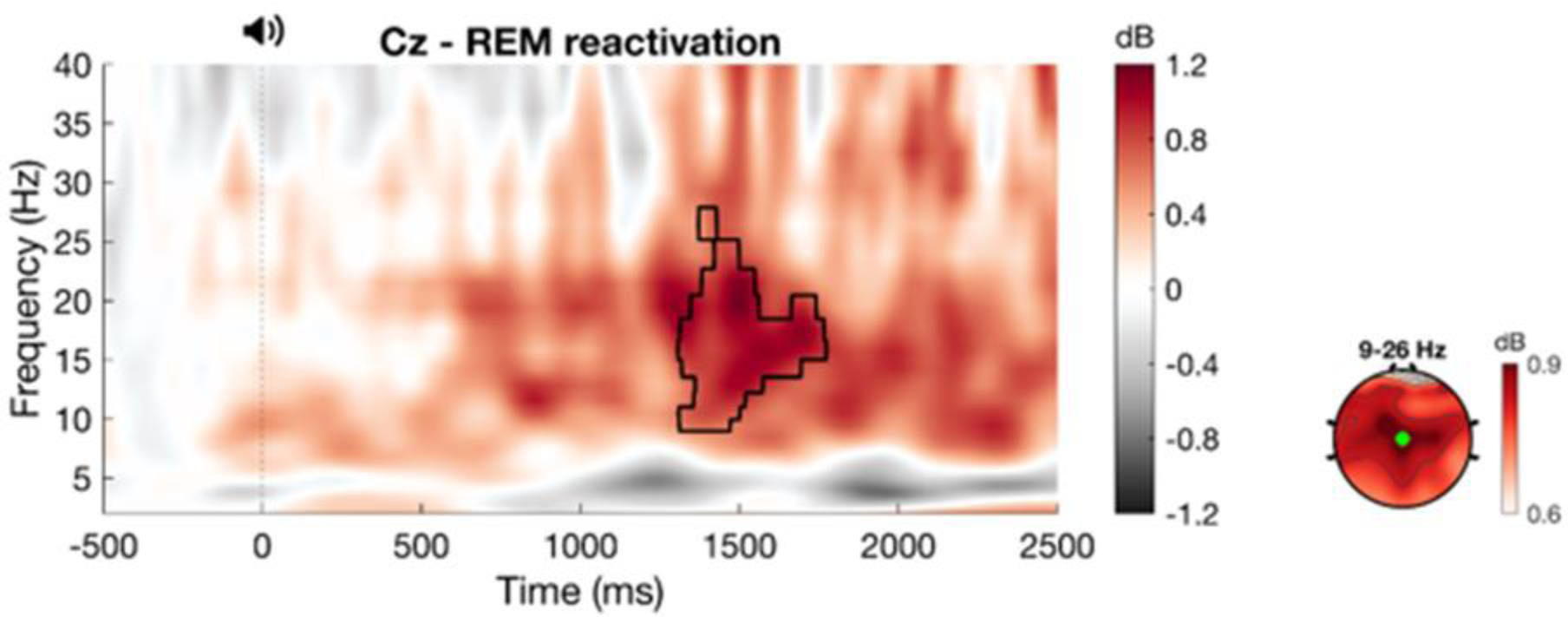
Effect of cueing during rapid eye movement sleep. Left: Time frequency response to emotional cue presentation during rapid eye movement sleep at electrode Cz. Time zero represents the initiation of sound presentation during sleep. Black contour line highlights the significant cluster of activation (*p* < .05, FDR adjusted). Right: Topographical visualizations of cluster-averaged activity.

No significant clusters emerged at the < .05 (FDR corrected) level in N-SWS (**Supplementary Figure S2).**

### 3.D Association of Sleep Fragmentation and Arousal with Memory

A recent study showed that participants who were awakened by reactivations showed memory impairment for cued items (Goldi and Rasch, 2019). We asked whether the memory impairment in E-REM group was associated with sleep disruptions, carrying out correlations between CB-T2 and arousal index (AI; number of arousals/minute; (Berry et al., 2012)), sleep fragmentation index (SFI; number of sleep stage transitions/minute; (Haba-Rubio et al., 2004)) and the duration of wake time after sleep onset (WASO). None of these revealed a significant association (AI: spearman’s r_s_ =-.29, p=.18; SFI: spearman’s r_s_ =.12, p=.58; WASO: spearman’s r_s_ =.26, p=.21). Because arousals could limit the number of reactivations, we also examined the correlation between CB-T2 and number of reactivations, but this also was not significant (spearman’s r_s_ =.10, p=.63). No significant associations were observed when the same analyses were repeated in the other reactivation groups.

Another recent TMR study found an inverse correlation of cueing benefit with cue-evoked power in beta band, and a strong trend for a similar correlation with power in alpha band (Whitmore et al., 2022). To examine if a similar mechanism explained the memory impairment for the reactivated items in E-REM and the negative cueing benefits in some participants in the other reactivation groups, we took the absolute difference in spectral power between the pre- and post-cue periods in the alpha (8-11 Hz) and beta (17-21 Hz) bands as an EEG-measure of cue-related arousal (Whitmore et al., 2022). While there were no correlations between these measures and the cueing benefit in the E-REM group, a significant negative correlation was observed between alpha power and CB-T2 (spearman’s r_s_ = -.53, p = .01) and CB-T3 (spearman’s r_s_ = -.47, p = .03) in E-SWS, similar to that seen by Whitmore et al. (2022). This suggests that higher cue-evoked alpha activity might contribute to the negative effect of SWS-TMR on emotional memory.

## 4. DISCUSSION

We applied TMR in SWS and REM to test the hypothesis that consolidation of emotional memories occurs primarily in REM. Contrary to our expectations, reactivation of emotional stimuli during REM led to poorer recall. A role for REM in memory decay was further supported by the association of higher %REM with reduced recall in the group that took a nap with no reactivation. On the other hand, the memory benefit of emotional memory reactivation during SWS was strongly correlated with the product of SWS and REM times (SWS-REM product), and in those with high SWS-REM product, reactivation during SWS significantly enhanced memory. Surprisingly, SWS-REM product was associated with enhanced memory for the reactivated and impaired memory for the non-reactivated items, suggesting a dual role for sleep in both strengthening and weakening memories. We also tested the hypothesis that emotional memories would be retained better than neutral ones after sleep. In support of this hypothesis, we found that reactivation in SWS led to larger memory benefit for emotional memories compared to neutral ones.

The decrease in recall after reactivation during REM was unexpected, yet it was consistent with what we observed in the nap only group (E-Nap), *i.e.,* more REM was associated with impaired recall. To the best of our knowledge, this is the first TMR study to show such an effect associated with REM. Our findings seemingly contradict the notion that REM sleep supports emotional memory consolidation. However, as reviewed recently (Davidson et al., 2021), evidence so far does not unequivocally support this notion. A role for REM in eliminating memories was first proposed by Crick and Mitchison (Crick and Mitchison, 1983). Later, different lines of research provided evidence suggesting that REM may be involved in selective removal of memory representations (Feld and Born, 2017; Poe, 2017; Langille, 2019), however, a direct link with forgetting in humans has been missing. While theta peak activity is associated with long-term potentiation in the hippocampus, several studies found that activity in the troughs causes depotentiation (Huerta and Lisman, 1995; Holscher et al., 1997). In one study, hippocampal place cells reversed their discharge phase to theta troughs, during REM once the exposed environment became familiar, suggesting that REM may serve to “refresh” synapses for future use in encoding new memories (Poe et al., 2000). Another study showed that firing rates of hippocampal neurons were decreased during REM and the decrease was correlated with REM theta (Grosmark et al., 2012). Furthermore, two recent studies showed direct evidence of synaptic pruning during REM after learning (Li et al., 2017; Zhou et al., 2020). Finally, direct behavioral evidence for REM related memory decay was found in a recent study (Izawa et al., 2019), which showed that a subset of melanin-concentrating-hormone-producing neurons were active specifically in REM and their activation was associated with impairment in memory. In light of this evidence, we can speculate that REM may have led to the weakening or elimination of synapses that represent emotional memories.

Memory transformation might also have contributed to the association of REM with memory impairment (possibly partially via the mechanisms summarized above), although we did not use any measures to examine this possibility. REM is particularly conducive to forming new associations and might have facilitated extraction of common elements across stimuli (Lewis, 2018), such as bodily harm, at the expense of information specific to individual stimulus (spatial location) (Langille, 2019). Another possibility is that we are seeing the effects of an “emotional memory trade-off”, characterized by sleep selectively enhancing emotional aspects of memory (picture content) while memory for less salient elements (spatial location) weakens (Denis et al., 2022). Finally, decline in memory might be related to the TMR method itself. In recent TMR studies, decreased recall was associated with cueing related increase in power in the alpha and beta bands (Whitmore et al., 2022) and memory for new vocabulary was impaired for the cued items, when the sound cues resulted in awakenings (Goldi and Rasch, 2019). We did not find any association of change in memory with arousals (AI or SI) but found increased alpha/beta activity in REM reactivation group (E-REM). However, this increase was not correlated with change in memory, and we observed that REM was associated with worse recall in a group with no reactivations, which makes it unlikely that results can be explained by only alpha/beta increase.

Furthermore, additional mechanisms can explain the cue-evoked activity we observed in the REM group. A recent study which carried out intracranial recordings found that anterior cingulate cortex (ACC) and dorsolateral prefrontal cortex (DLPFC) showed bursts of beta activity during REM, in addition to the expected theta activity (Vijayan et al., 2017). Furthermore, another study which used auditory stimulation to enhance theta activity, observed increased power in the alpha/beta band similar to our study (Harrington et al., 2021). While we did not observe an increase in theta oscillations, increased beta activity may be related to activation in regions involved in emotional processing, such as ACC.

The strong correlation of cueing benefit with the product of SWS and REM times in the E-SWS group suggests a complementary role for these sleep stages. This finding is in agreement with accounts that propose that both sleep stages are required for some forms of sleep-dependent memory consolidation (Giuditta et al., 1995; Ficca and Salzarulo, 2004; Diekelmann and Born, 2010; Walker and Stickgold, 2010; Langille, 2019) and the expanding literature which provides supporting evidence in humans. Studies using a visual discrimination task found that improvement was correlated with the product of times spent in SWS and REM (Stickgold et al., 2000), largest after a full night of sleep compared to early or late night-half sleep (Gais et al., 2000), and present after a nap only if it contained both NREM and REMs (Mednick et al., 2003). In studies that used verbal memory tasks, recall was positively correlated with the duration of NREM/REM cycles (Mazzoni et al., 1999) and was impaired when sleep cycles were interrupted, but not after sleep fragmentation with intact sleep cycles (Ficca et al., 2000). Finally, similar to our study, several TMR studies that carried out reactivations in SWS found that REM or both REM and SWS were associated with outcomes. REM duration was the only measure correlated with enhanced memory for the meaning of novel words (Batterink et al., 2017) and lexical integration (Tamminen et al., 2017) after TMR. In another study, TMR related reduction in social bias was correlated with %SWS × %REM (Hu et al., 2015). Besides behavioral effects, other TMR studies with reactivation in SWS, found neurobiological alterations associated with REM (Cairney et al., 2015, Cousins et al., 2016). In summary, there is now a substantial body of evidence which suggests that memory consolidation is attained via processes that take place across NREM and REM sleep. Further studies are needed to directly examine the dynamics of this coordination, such as how brain oscillations interact across sleep stages.

The product of SWS and REM times was associated with enhanced memory for reactivated items and impaired memory for items that were not reactivated. This suggests that sleep provides memory advantage by not only increasing the retention (or reducing the forgetting) of select memories, but by also enhancing the forgetting of irrelevant ones, and that forgetting occurs across SWS and REM. These findings are consistent with models which propose that active forgetting complements memory consolidation and implicate both SWS and REM in this process (Tononi and Cirelli 2014, Feld and Born 2017). Indeed, in addition to the REM mechanisms that may potentially facilitate memory impairment summarized above, recent evidence indicates that SWS may also lead to weakening of memory representations (Colgin et al., 2004, Tononi and Cirelli 2014, Andrillon et al., 2017, Gonzalez-Rueda et al., 2018, Norimoto et al., 2018, Kim et al., 2019). Moreover, a recent study showed that hippocampal firing rate decrease during sleep was predicted by sleep spindles and sharp wave ripples, but was not evident until the following REM, suggesting that coordinated activity of SWS and REM might be involved in memory impairment. In our study, TMR may have protected the cued items from decay.

Cueing emotional stimuli during SWS generated EEG responses in the delta/theta band and then at spindle frequencies, similar to previous studies (Lehmann et al., 2016; Forcato et al., 2020; Schechtman et al., 2021). Notably, valence of the sound cues was correlated with activity increase in both frequency ranges, suggesting that they contribute to emotional processing during sleep. That spindle activity was modulated by emotional valence was supported by the additional finding that power in the spindle band was the only difference in cue-evoked activity between E-SWS and N-SWS groups, with significantly higher spindle power in E-SWS. A role for spindles in emotional processing has been suggested in recent studies that showed an association of spindle activity with enhanced consolidation of emotional memories (Cairney et al., 2014a; Lehmann et al., 2016; Alger et al., 2018) and with reduction in reactivity to negative memories after sleep (Azza et al., 2022). In addition to correlations, another study (Kaestner et al., 2013) found that increasing spindle activity pharmacologically significantly improved memory for negative and high-arousal items. Aside from experimental studies, spindles are also implicated in the pathophysiology of psychiatric disorders including social anxiety disorder (Wilhelm et al., 2017) and major depressive disorder (Lopez et al., 2010; Plante et al., 2013; Nishida et al., 2014; Sesso et al., 2017). Furthermore, altered spindle activity is associated with specific emotional symptoms, including worry (Hamann et al., 2019), internalizing problems (Kathrin et al., 2021), intrusive memories (van der Heijden et al., 2022), and frequent nightmares (Picard-Deland et al., 2018). Our results lend further support to the burgeoning evidence suggesting that spindles may be involved in emotional processing. There was no cue-evoked activity increase in N-SWS, which could be due to neutral sounds not being sufficiently discernible or failure to establish strong sound-picture associations.

Taken together, our findings suggest that i. emotional memories decay during REM, unless they were reactivated during prior SWS; ii. forgetting is a complementary process in the consolidation of emotional memories; and iii. both SWS and REM are required to preserve relevant memories and eliminate irrelevant ones. We also show that in addition to delta/theta activity, sleep spindles are modulated by emotional charge of memories. Finally, emotional memories were retained better than neutral ones as a result of reactivation in SWS. However, we cannot rule out the possibility that this was simply due to failure to induce any effect on memory in the group that was exposed to neutral items.

The authors declare no competing financial interests.

## Acknowledgments

This work was funded by NIH grant MH048832 to Dr. Stickgold.

## Supplementary Material

### Pilot Study

The pilot study included 4 phases carried out in two identical rounds in two different samples. In the first three phases, participants rated the stimuli for valence and arousal. They first rated sounds-only (phase 1), then sound-picture pairs (phase 2), and then the sounds-only again (phase 3). Repeating the sounds only ratings after sound-picture pairs allowed us to see if the ratings changed after participants were exposed to the corresponding pictures. Ratings were carried out by sliding a marker across a happy-unhappy (for valence) or excited-calm (for arousal) mannequins displayed on the screen, after being exposed to each stimulus. In the fourth phase, participants were asked to match the sounds they heard with their corresponding images. The first round included 49 emotional and 49 neutral pictures. One more sound-picture pair was added to both the neutral and emotional stimulus sets after the first round, making the totals 50 in both sets. 20 healthy individuals, 18-25 years old, who were not later included in the main study, participated. Participants reported no history of sleep abnormalities or psychiatric disorders. In each round, they were assigned to either a neutral or emotional stimulus set.

In the first round, valence ratings were significantly different between neutral and emotional sets in all phases (p<.001), with emotional items rated as more negative. Arousal ratings were different only for sound-picture pairs (p<.001), with emotional items rated as more arousing, but not for sounds-only in phase 1 or phase 3 (p=.12 and p=.38, respectively). Because one of the items in the neutral set was rated as highly negative, it was moved to the emotional set for the second round. Another item in the emotional set was rated similar to neutral items, so it was removed after the first round. In the second round, similar to the first round, emotional items were rated more negative in all phases (p=.01, p<.001 and p<.001, respectively). Sounds-only in phase 3 were rated more arousing (p=.002), however, there were no differences in arousal ratings for sounds only in phase 1 or for sound-picture pairs in phase 2 (p=.36 and p=.09, respectively).

**Table S1.**
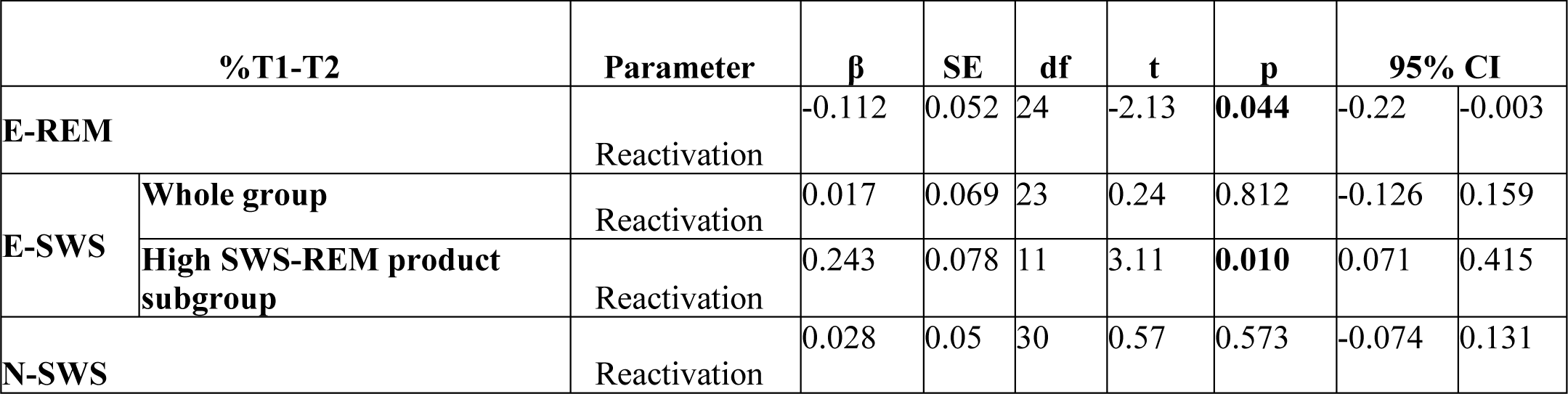
Linear mixed-effects models in separate groups for %T1-T2.

**Table S2.**
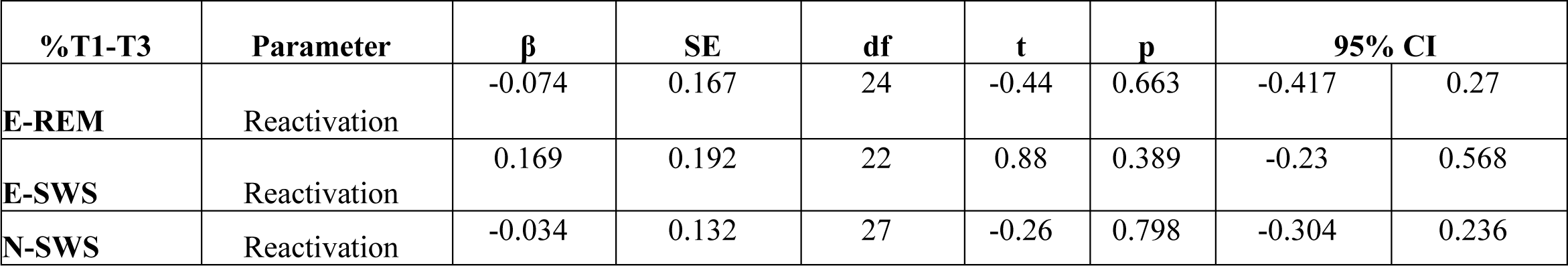
Linear mixed-effects models in separate groups for %T1-T3.

**Table S3.**
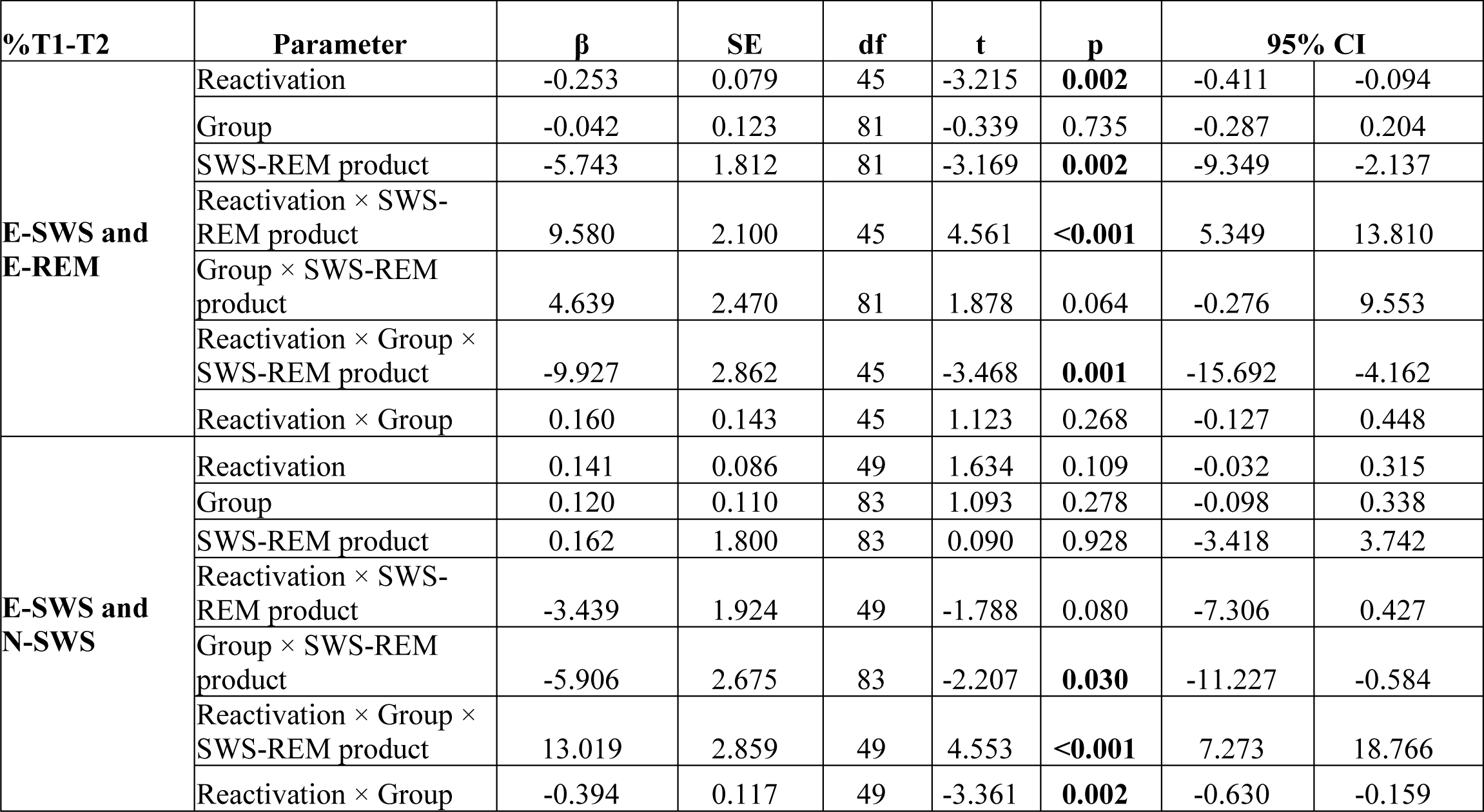
Linear mixed-effects models which compare the effect of reactivation on %T1-T2 in E-SWS vs. E-REM, and E-SWS vs. N-SWS.

| %T1-T2 | Parameter | $\beta$ | SE | df | t | p | 95% CI | |
| --- | --- | --- | --- | --- | --- | --- | --- | --- |
| **E-SWS and E-REM** | Reactivation | -0.253 | 0.079 | 45 | -3.215 | **0.002** | -0.411 | -0.094 |
|  | Group | -0.042 | 0.123 | 81 | -0.339 | 0.735 | -0.287 | 0.204 |
|  | SWS-REM product | -5.743 | 1.812 | 81 | -3.169 | **0.002** | -9.349 | -2.137 |
| | Reactivation × SWS-REM product | 9.580 | 2.100 | 45 | 4.561 | **<0.001** | 5.349 | 13.810 |
| | Group × SWS-REM product | 4.639 | 2.470 | 81 | 1.878 | 0.064 | -0.276 | 9.553 |
| | Reactivation × Group × SWS-REM product | -9.927 | 2.862 | 45 | -3.468 | **0.001** | -15.692 | -4.162 |
| | Reactivation × Group | 0.160 | 0.143 | 45 | 1.123 | 0.268 | -0.127 | 0.448 |
| **E-SWS and N-SWS** | Reactivation | 0.141 | 0.086 | 49 | 1.634 | 0.109 | -0.032 | 0.315 |
|  | Group | 0.120 | 0.110 | 83 | 1.093 | 0.278 | -0.098 | 0.338 |
|  | SWS-REM product | 0.162 | 1.800 | 83 | 0.090 | 0.928 | -3.418 | 3.742 |
| | Reactivation × SWS-REM product | -3.439 | 1.924 | 49 | -1.788 | 0.080 | -7.306 | 0.427 |
| | Group × SWS-REM product | -5.906 | 2.675 | 83 | -2.207 | **0.030** | -11.227 | -0.584 |
| | Reactivation × Group × SWS-REM product | 13.019 | 2.859 | 49 | 4.553 | **<0.001** | 7.273 | 18.766 |
| | Reactivation × Group | -0.394 | 0.117 | 49 | -3.361 | **0.002** | -0.630 | -0.159 |

**Table S4.**
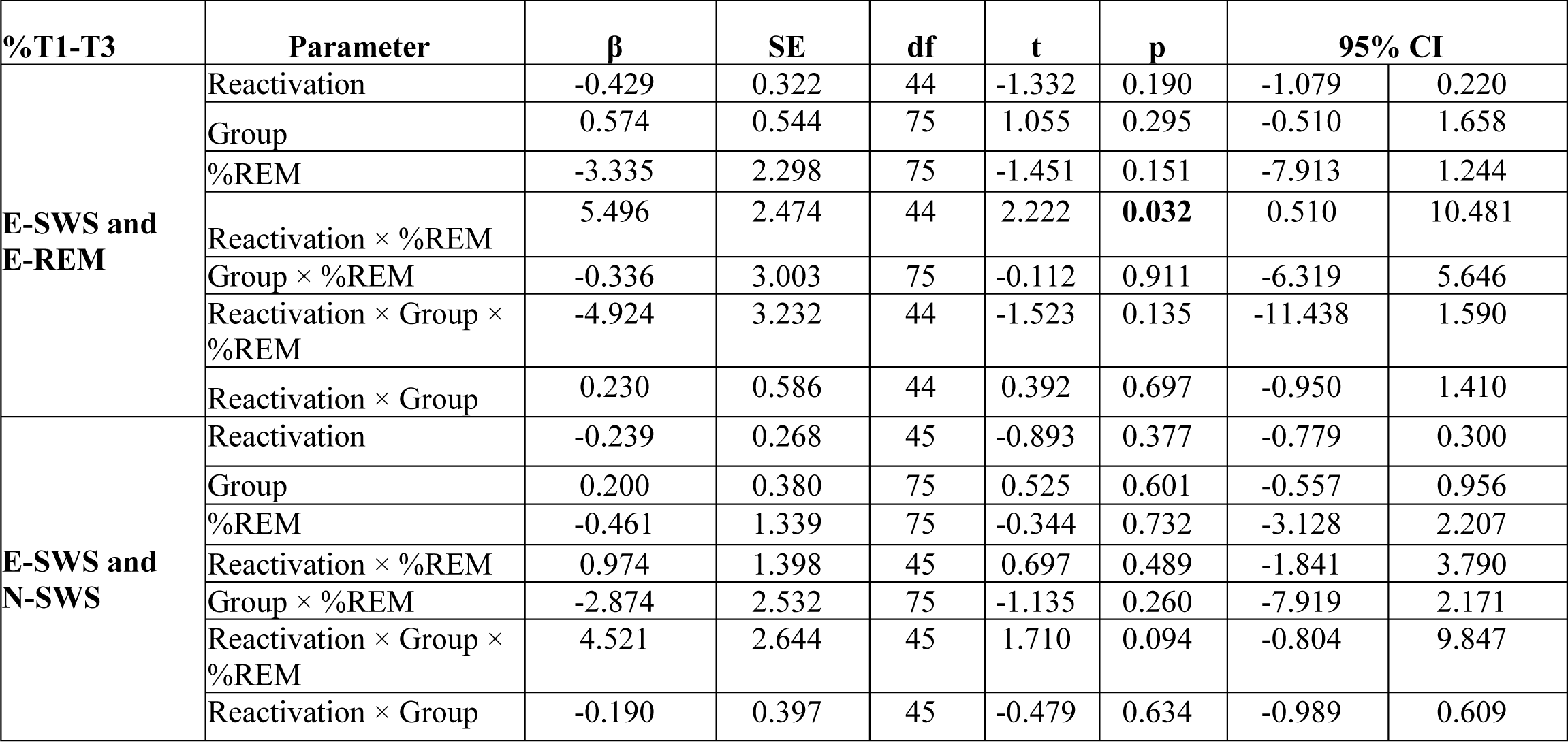
Linear mixed-effects models which compare the effect of reactivation on %T1-T3 in E-SWS vs. E-REM, and E-SWS vs. N-SWS.

| %T1-T3 | Parameter | $\beta$ | SE | df | t | p | 95% CI | |
| --- | --- | --- | --- | --- | --- | --- | --- | --- |
| **E-SWS and E-REM** | Reactivation | -0.429 | 0.322 | 44 | -1.332 | 0.190 | -1.079 | 0.220 |
|  | Group | 0.574 | 0.544 | 75 | 1.055 | 0.295 | -0.510 | 1.658 |
|  | %REM | -3.335 | 2.298 | 75 | -1.451 | 0.151 | -7.913 | 1.244 |
| | Reactivation × %REM | 5.496 | 2.474 | 44 | 2.222 | **0.032** | 0.510 | 10.481 |
| | Group × %REM | -0.336 | 3.003 | 75 | -0.112 | 0.911 | -6.319 | 5.646 |
| | Reactivation × Group × %REM | -4.924 | 3.232 | 44 | -1.523 | 0.135 | -11.438 | 1.590 |
| | Reactivation × Group | 0.230 | 0.586 | 44 | 0.392 | 0.697 | -0.950 | 1.410 |
| **E-SWS and N-SWS** | Reactivation | -0.239 | 0.268 | 45 | -0.893 | 0.377 | -0.779 | 0.300 |
|  | Group | 0.200 | 0.380 | 75 | 0.525 | 0.601 | -0.557 | 0.956 |
|  | %REM | -0.461 | 1.339 | 75 | -0.344 | 0.732 | -3.128 | 2.207 |
| | Reactivation × %REM | 0.974 | 1.398 | 45 | 0.697 | 0.489 | -1.841 | 3.790 |
| | Group × %REM | -2.874 | 2.532 | 75 | -1.135 | 0.260 | -7.919 | 2.171 |
| | Reactivation × Group × %REM | 4.521 | 2.644 | 45 | 1.710 | 0.094 | -0.804 | 9.847 |
| | Reactivation × Group | -0.190 | 0.397 | 45 | -0.479 | 0.634 | -0.989 | 0.609 |

**Table S5.**
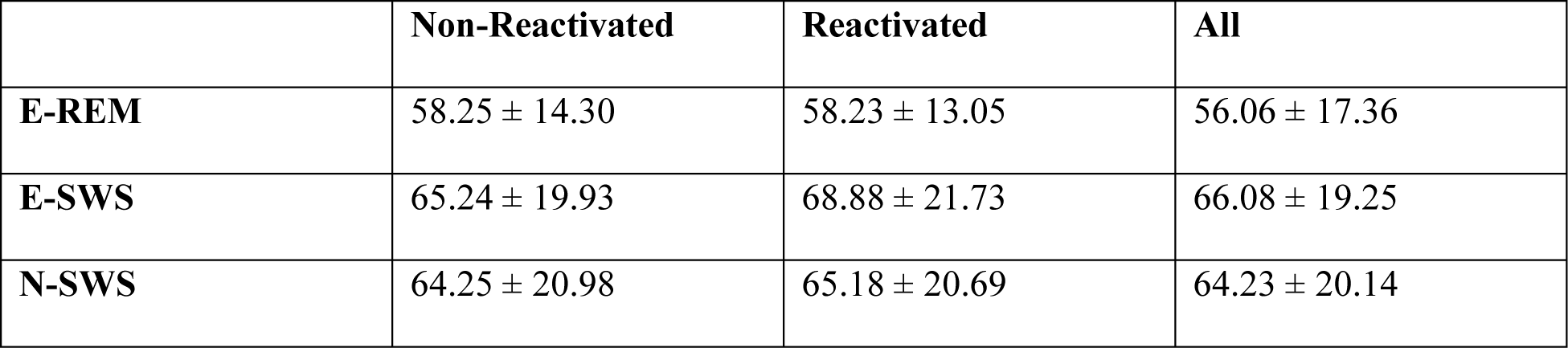
Recall at T1 (baseline memory). There was no difference between groups, or between non-reactivated and reactivated items within any of the groups. See “Results” section for the statistical analyses. Data is displayed as mean ± standard deviation.

**Table S6.**
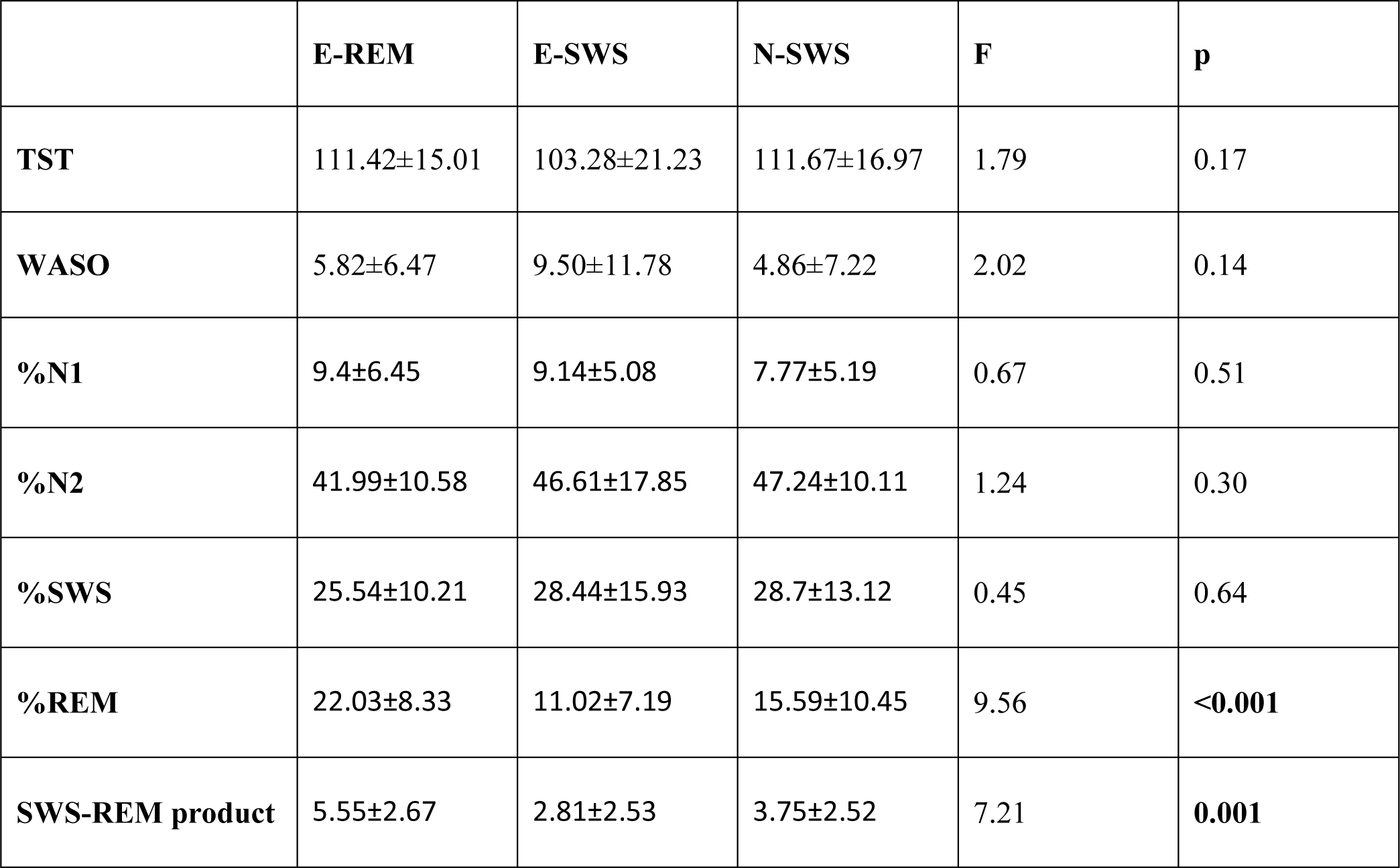
Sleep variables between reactivation groups are compared using one-way MANOVA. Post-hoc pairwise comparisons showed that E-REM group had higher %REM and %SWS × %REM compared to other groups. Data is displayed as mean ± standard deviation.

**Figure S1.**
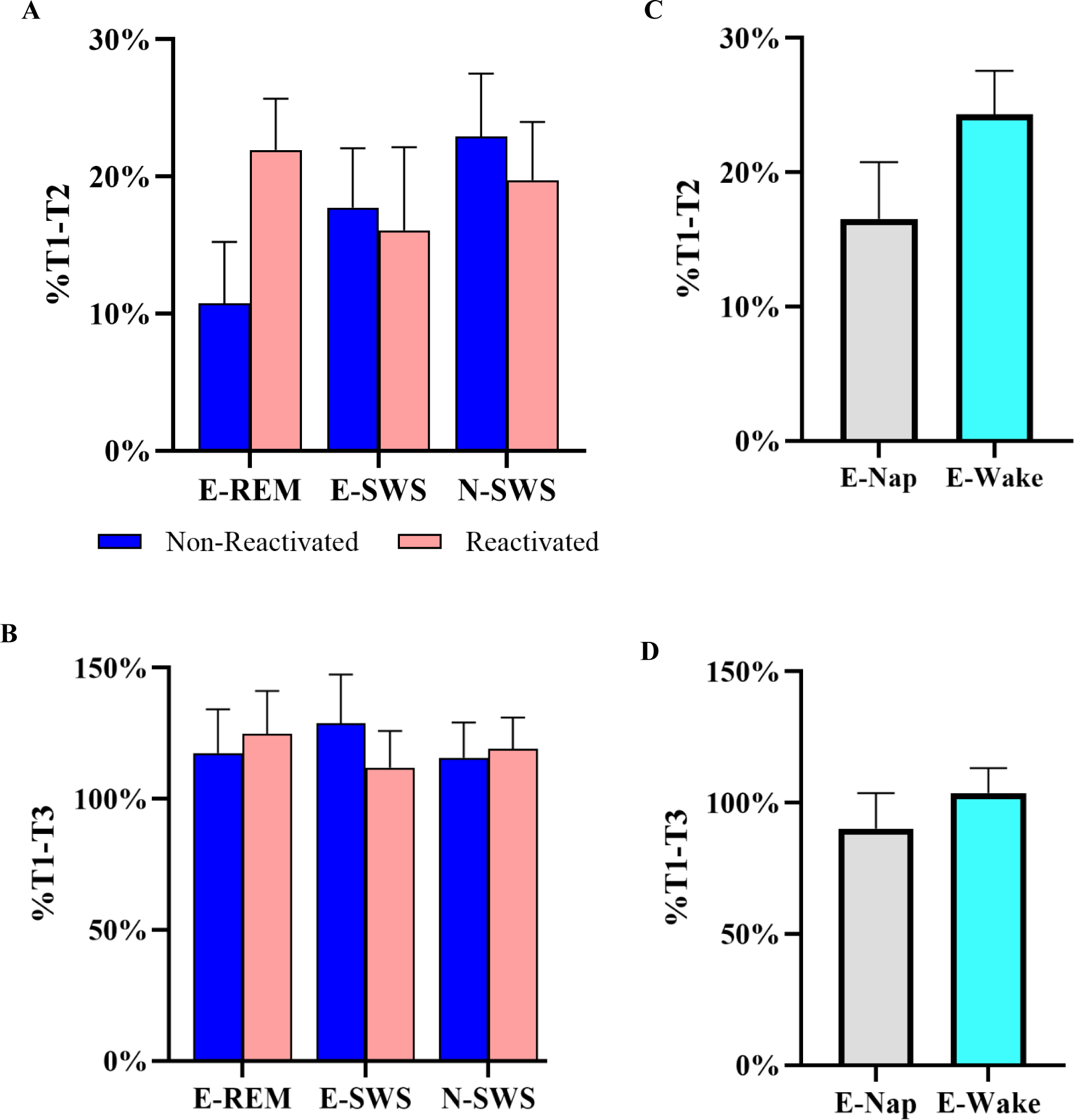
Observed means of %change in error from baseline (T1) to retest (T2; %T1-T2) and to delayed retest (T3; %T1-T3), in reactivation groups (A and B), and in nap and wake groups (C and D). Error bars indicate the standard error.

**Figure S2.**
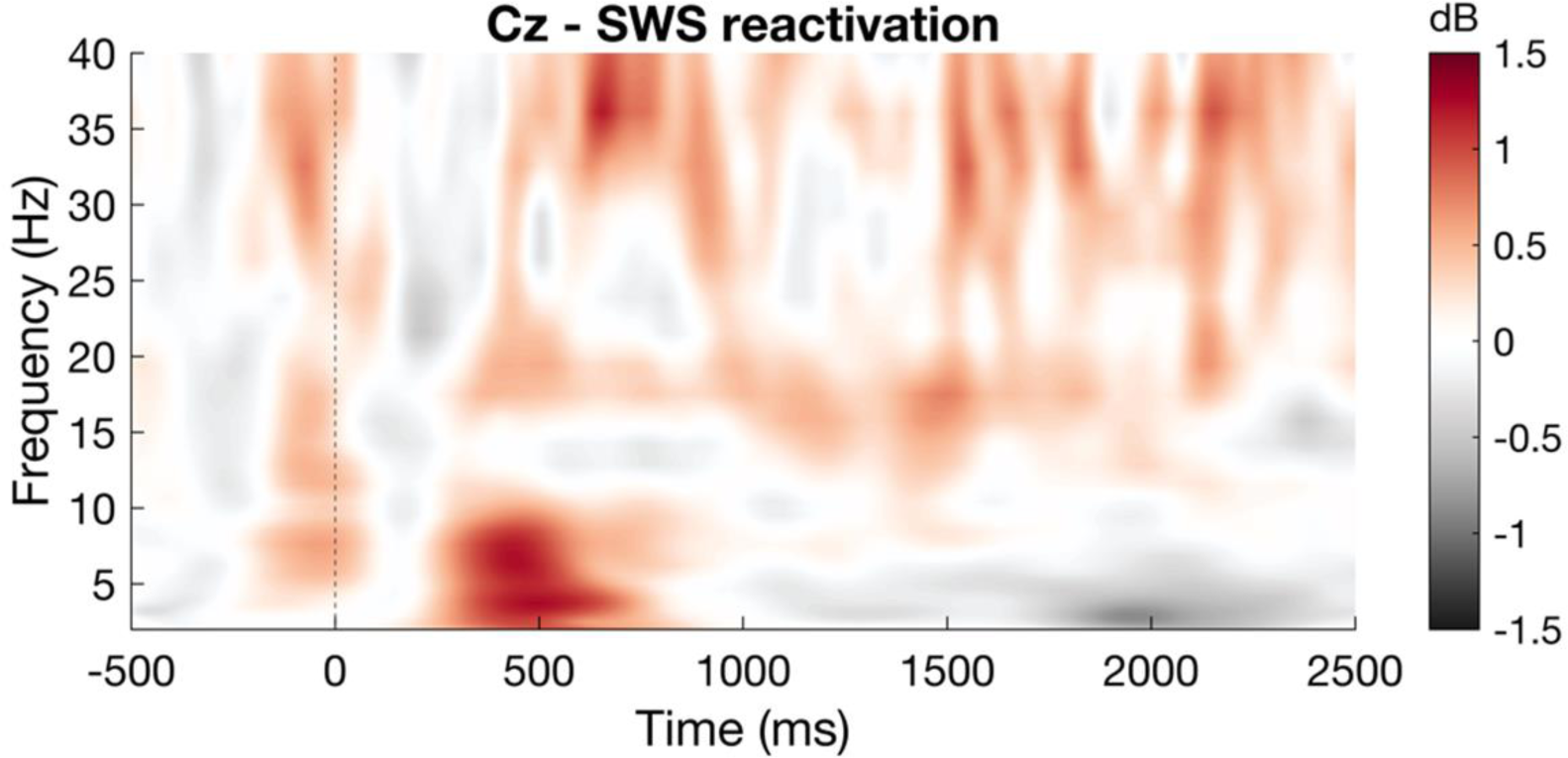
Time frequency response to neutral cue presentation during slow wave sleep at electrode Cz. Time zero represents the initiation of sound presentation during sleep. No significant clusters of activity emerged.

## Notes

### Competing Interest Statement

The authors have declared no competing interest.

### Summary of Updates

We discovered errors with the statistical approach and behavioral data. Additionally, we revised the presentation of our results, to enhance their clarity and comprehensibility.

