## Supplementary Materials for "Emotional memories are enhanced when reactivated in slow-wave sleep but impaired in REM"

| %T1-T2 | | Parameter | β | SE | df | t | p | 95% CI | |
|---|---|---|---|---|---|---|---|---|---|
| **E-REM** | | Reactivation | -0.112 | 0.052 | 24 | -2.13 | **0.044** | -0.22 | -0.003 |
| **E-SWS** | **Whole group** | Reactivation | 0.017 | 0.069 | 23 | 0.24 | 0.812 | -0.126 | 0.159 |
| | **High SWS-REM product subgroup** | Reactivation | 0.243 | 0.078 | 11 | 3.11 | **0.010** | 0.071 | 0.415 |
| **N-SWS** | | Reactivation | 0.028 | 0.05 | 30 | 0.57 | 0.573 | -0.074 | 0.131 |

**Table S1.** Linear mixed-effects models in separate groups for %T1-T2.

| %T1-T3 | Parameter | β | SE | df | t | p | 95% CI | |
|---|---|---|---|---|---|---|---|---|
| **E-REM** | Reactivation | -0.074 | 0.167 | 24 | -0.44 | 0.663 | -0.417 | 0.27 |
| **E-SWS** | Reactivation | 0.169 | 0.192 | 22 | 0.88 | 0.389 | -0.23 | 0.568 |
| **N-SWS** | Reactivation | -0.034 | 0.132 | 27 | -0.26 | 0.798 | -0.304 | 0.236 |

**Table S2.** Linear mixed-effects models in separate groups for %T1-T3.

| %T1-T2 | Parameter | β | SE | df | t | p | 95% CI | |
|---|---|---|---|---|---|---|---|---|
| **E-SWS and E-REM** | Reactivation | -0.253 | 0.079 | 45 | -3.215 | **0.002** | -0.411 | -0.094 |
| | Group | -0.042 | 0.123 | 81 | -0.339 | 0.735 | -0.287 | 0.204 |
| | SWS-REM product | -5.743 | 1.812 | 81 | -3.169 | **0.002** | -9.349 | -2.137 |
| | Reactivation × SWS-REM product | 9.580 | 2.100 | 45 | 4.561 | **<0.001** | 5.349 | 13.810 |
| | Group × SWS-REM product | 4.639 | 2.470 | 81 | 1.878 | 0.064 | -0.276 | 9.553 |
| | Reactivation × Group × SWS-REM product | -9.927 | 2.862 | 45 | -3.468 | **0.001** | -15.692 | -4.162 |
| | Reactivation × Group | 0.160 | 0.143 | 45 | 1.123 | 0.268 | -0.127 | 0.448 |
| **E-SWS and N-SWS** | Reactivation | 0.141 | 0.086 | 49 | 1.634 | 0.109 | -0.032 | 0.315 |
| | Group | 0.120 | 0.110 | 83 | 1.093 | 0.278 | -0.098 | 0.338 |
| | SWS-REM product | 0.162 | 1.800 | 83 | 0.090 | 0.928 | -3.418 | 3.742 |
| | Reactivation × SWS-REM product | -3.439 | 1.924 | 49 | -1.788 | 0.080 | -7.306 | 0.427 |
| | Group × SWS-REM product | -5.906 | 2.675 | 83 | -2.207 | **0.030** | -11.227 | -0.584 |
| | Reactivation × Group × SWS-REM product | 13.019 | 2.859 | 49 | 4.553 | **<0.001** | 7.273 | 18.766 |
| | Reactivation × Group | -0.394 | 0.117 | 49 | -3.361 | **0.002** | -0.630 | -0.159 |

**Table S3.** Linear mixed-effects models which compare the effect of reactivation on %T1-T2 in E-SWS vs. E-REM, and E-SWS vs. N-SWS.

| %T1-T3 | Parameter | β | SE | df | t | p | 95% CI | |
|---|---|---|---|---|---|---|---|---|
| E-SWS and E-REM | Reactivation | -0.429 | 0.322 | 44 | -1.332 | 0.190 | -1.079 | 0.220 |
| | Group | 0.574 | 0.544 | 75 | 1.055 | 0.295 | -0.510 | 1.658 |
| | %REM | -3.335 | 2.298 | 75 | -1.451 | 0.151 | -7.913 | 1.244 |
| | Reactivation × %REM | 5.496 | 2.474 | 44 | 2.222 | **0.032** | 0.510 | 10.481 |
| | Group × %REM | -0.336 | 3.003 | 75 | -0.112 | 0.911 | -6.319 | 5.646 |
| | Reactivation × Group × %REM | -4.924 | 3.232 | 44 | -1.523 | 0.135 | -11.438 | 1.590 |
| | Reactivation × Group | 0.230 | 0.586 | 44 | 0.392 | 0.697 | -0.950 | 1.410 |
| E-SWS and N-SWS | Reactivation | -0.239 | 0.268 | 45 | -0.893 | 0.377 | -0.779 | 0.300 |
| | Group | 0.200 | 0.380 | 75 | 0.525 | 0.601 | -0.557 | 0.956 |
| | %REM | -0.461 | 1.339 | 75 | -0.344 | 0.732 | -3.128 | 2.207 |
| | Reactivation × %REM | 0.974 | 1.398 | 45 | 0.697 | 0.489 | -1.841 | 3.790 |
| | Group × %REM | -2.874 | 2.532 | 75 | -1.135 | 0.260 | -7.919 | 2.171 |
| | Reactivation × Group × %REM | 4.521 | 2.644 | 45 | 1.710 | 0.094 | -0.804 | 9.847 |
| | Reactivation × Group | -0.190 | 0.397 | 45 | -0.479 | 0.634 | -0.989 | 0.609 |

**Table S4.** Linear mixed-effects models which compare the effect of reactivation on %T1-T3 in E-SWS vs. E-REM, and E-SWS vs. N-SWS.

| | Non-Reactivated | Reactivated | All |
|---|---|---|---|
| **E-REM** | 58.25 ± 14.30 | 58.23 ± 13.05 | 56.06 ± 17.36 |
| **E-SWS** | 65.24 ± 19.93 | 68.88 ± 21.73 | 66.08 ± 19.25 |
| **N-SWS** | 64.25 ± 20.98 | 65.18 ± 20.69 | 64.23 ± 20.14 |

**Table S5.** Recall at T1 (baseline memory). There was no difference between groups, or between non-reactivated and reactivated items within any of the groups. See "Results" section for the statistical analyses. Data is displayed as mean ± standard deviation.

| | E-REM | E-SWS | N-SWS | F | p |
|---|---|---|---|---|---|
| **TST** | 111.42±15.01 | 103.28±21.23 | 111.67±16.97 | 1.79 | 0.17 |
| **WASO** | 5.82±6.47 | 9.50±11.78 | 4.86±7.22 | 2.02 | 0.14 |
| **%N1** | 9.4±6.45 | 9.14±5.08 | 7.77±5.19 | 0.67 | 0.51 |
| **%N2** | 41.99±10.58 | 46.61±17.85 | 47.24±10.11 | 1.24 | 0.30 |
| **%SWS** | 25.54±10.21 | 28.44±15.93 | 28.7±13.12 | 0.45 | 0.64 |
| **%REM** | 22.03±8.33 | 11.02±7.19 | 15.59±10.45 | 9.56 | **<0.001** |
| **SWS-REM product** | 5.55±2.67 | 2.81±2.53 | 3.75±2.52 | 7.21 | **0.001** |

**Table S6.** Sleep variables between reactivation groups are compared using one-way MANOVA. Post-hoc pairwise comparisons showed that E-REM group had higher %REM and %SWS × %REM compared to other groups. Data is displayed as mean ± standard deviation.

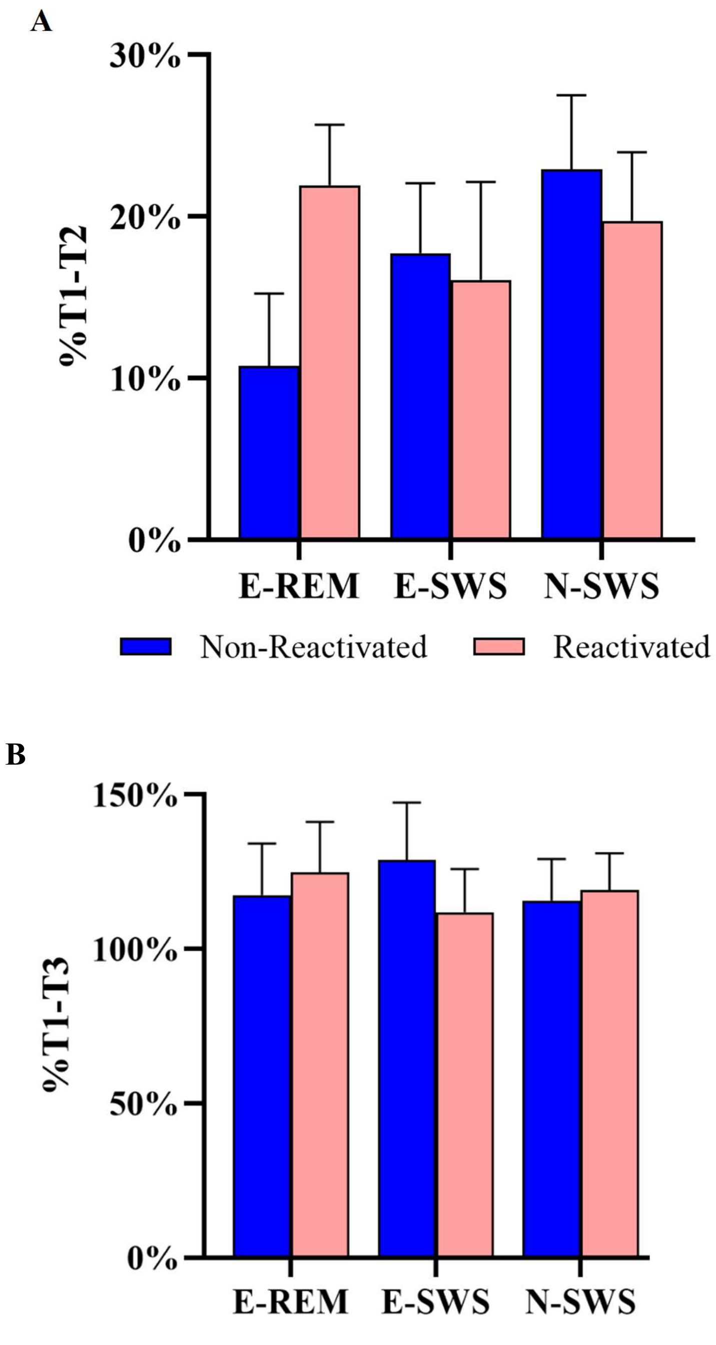


**Figure S1.** Observed means of %change in error from baseline (T1) to retest (T2; %T1-T2) and to delayed retest (T3; %T1-T3), in reactivation groups (A and B), and in nap and wake groups (C and D). Error bars indicate the standard error.

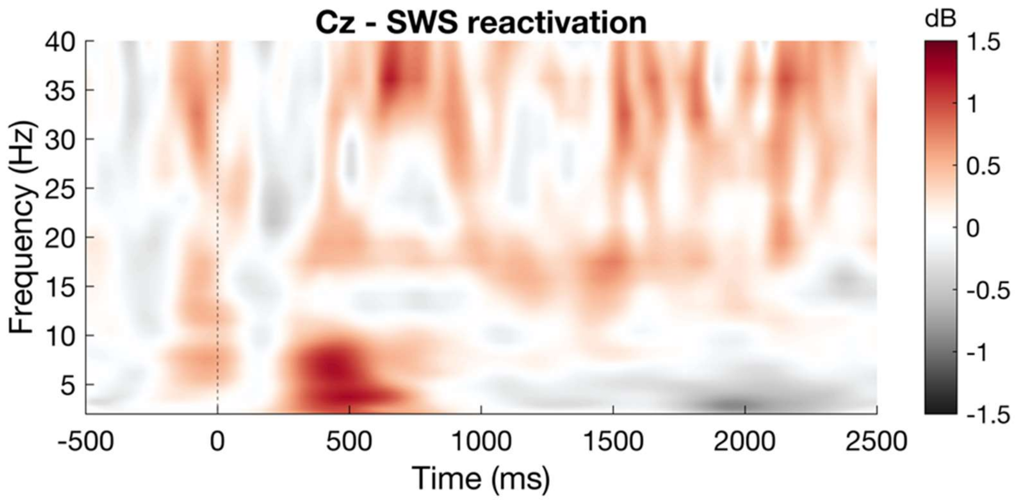


**Figure S2.** Time frequency response to neutral cue presentation during slow wave sleep at electrode Cz. Time zero represents the initiation of sound presentation during sleep. No significant clusters of activity emerged.
